# MELD-DNA: A new tool for capturing protein-DNA binding

**DOI:** 10.1101/2021.06.24.449809

**Authors:** Antonio Bauzá, Alberto Pérez

## Abstract

Herein we present MELD-DNA, a novel computational approach to address the problem of protein-DNA structure prediction. This method addresses well-known issues hampering current computational approaches to bridge the gap between structural and sequence knowledge, such as large conformational changes in DNA and highly charged electrostatic interaction during binding. MELD-DNA is able to: i) sample multiple binding modes, ii) identify the preferred binding mode from the ensembles, and iii) provide qualitative binding preferences between DNA sequences. We expect the results presented herein will have impact in the field of biophysics (through new software development), structural biology (by complementing DNA structural databases) and supramolecular chemistry (by bringing new insights into protein-DNA interactions).

---

According to the Nucleic Acid Database (NDB),^[1]^ 5600 protein-DNA structures are deposited in the Protein Data Bank (PDB),^[2]^ which represents a small fraction compared to our protein structural knowledge and the amount of DNA binding protein families known to date. Protein-DNA interactions play a crucial role in biology^[3]^ (e.g., in the storage, repair, expression and regulation of genetic information^[4,5][6]^) as well as in other areas of research (e. g. nanomedicine^[7]^ transition metal chemistry^[8]^ or even clinical diagnosis^[9]^). Whilst proteins that bind other proteins possess a limited number of binding partners, a protein that binds DNA is able to bind many sequences with high affinity. Many Protein-DNA interactions take place through non-specific sites like the phosphate backbone, driven by charge complementarity, resulting in promiscuous binding. Despite of this, the specificity^[10]^ (or relative binding affinity) to different sequences can span several orders of magnitude, leading to a preferential binding for certain sequences (defined as consensus sequences).

Protein-DNA recognition is generally driven by two mechanisms: base readout and shape readout^[11]^ The former mainly refers to interactions between protein and DNA and is generally believed to be responsible for determining the binding mode. On the other hand, the shape readout considers the ability of a certain DNA sequence to adopt a particular conformation needed for binding. Shape readout is believed to be important to modulate specificity – which DNA sequences will bind more favorably to a particular DNA binding protein. The relative importance of the base and shape readout mechanisms for DNA recognition varies from protein to protein, but there is a lack of quantitative methods able to predict the balance between the two^[12,13]^ In this regard, several methods have tried to quantify the balance of each or used coarse representations^[14]^ of the shape readout to predict sequences that can adopt certain conformations.^[15-18]^ Hence, a deeper understanding of the balance between the two mechanisms would certainly result in more accurate physical models that can in turn be used to grand challenges like the gene regulation problem: how and where transcription factor proteins bind along the genome^[19,20]^ We thus need tools that can predict binding and/or specificity, in an effort to diminish the sparsity in structural databases.

In this context, docking is the most efficient technique for predicting macromolecular binding.^[21]^ It is especially useful for the virtual screening of small molecules libraries in the early stages of drug discovery^[22]^ and has seen significant improvements for the study of protein-protein interactions thanks to community efforts like the Critical Assessment of PRediction of Interactions (CAPRI)^[23]^ However, current docking methods are not well suited to tackle the protein-DNA problem, which implies two major challenges; i) sampling large DNA conformational changes upon binding and ii) scoring the predictions due to the highly charged electrostatic interaction between proteins and DNA^[24]^ Consequently, the protein-DNA docking field has advanced at a slower pace than protein-protein docking. In this regard, computational efforts have been carried out by the HADDOCK^[25,26]^ team by combining ambiguous data with DNA flexibility as well as in compiling benchmark sets of different difficulty for assessing protein-DNA docking performance^[27]^

Predicting specificity based on the consensus binding mode is also a challenge for protein-DNA systems. Alchemical free energy methods based on Molecular Dynamics simulations (MD) are the gold standard for predicting relative binding affinities given a known binding mode^[28]^ achieving an unprecedented level of accuracy for predicting relative and absolute binding free energies in small molecules^[29]^ However, they present strong limitations when dealing with large charge variations and with flexible complexes where several binding modes can contribute to the binding free energy^[28]^ While alchemical free energy calculations have been successful in studying the specificity for some protein-DNA systems^[30,31]^ a recent systematic study^[32]^ on protein-DNA complexes points to deficiencies of these approaches arising from either phase space overlap or force field issues.

Bearing this in mind, we herein propose a novel computational methodology, named MELD-DNA, where we combine a new implicit solvent^[33]^ and force field^[34]^ to identify protein-nucleic acid structures with the MELD^[35]^ (Modeling Employing Limited Data) Bayesian inference approach. MELD i) is based on MD techniques, which naturally account for sequence-dependent DNA flexibility, ii) uses replica exchange (RE) approaches for enhanced sampling of binding/unbinding events, and iii) benefits from generic and ambiguous information to guide the sampling (see Computational Methods in SI). MELD has already achieved success in predicting protein structures^[36]^ as well as in protein binding to small molecules^[37]^ peptides^[38]^ and other proteins^[39]^ Herein we have applied this methodology to three systems involving different levels of DNA deformation (see Fig. 1a and SI): low (basic Leucine Zipper Domain, bZip), medium (P22-C2 repressor) and large (TATA box). MELD-DNA successfully: i) samples multiple binding modes, ii) identifies the experimental binding mode through clustering of the ensembles and iii) is sensitive to DNA sequences and conformations. As a general procedure, we have performed simulations in which the DNA conformation is either free to explore bound/B-DNA conformations or restrained to the bound conformation to differentiate between sequence and structural preferences (see SI for details and Fig. 1b). We further tested the value of competitive binding simulations for qualitative prediction of relative binding affinities.

**Figure 1.**
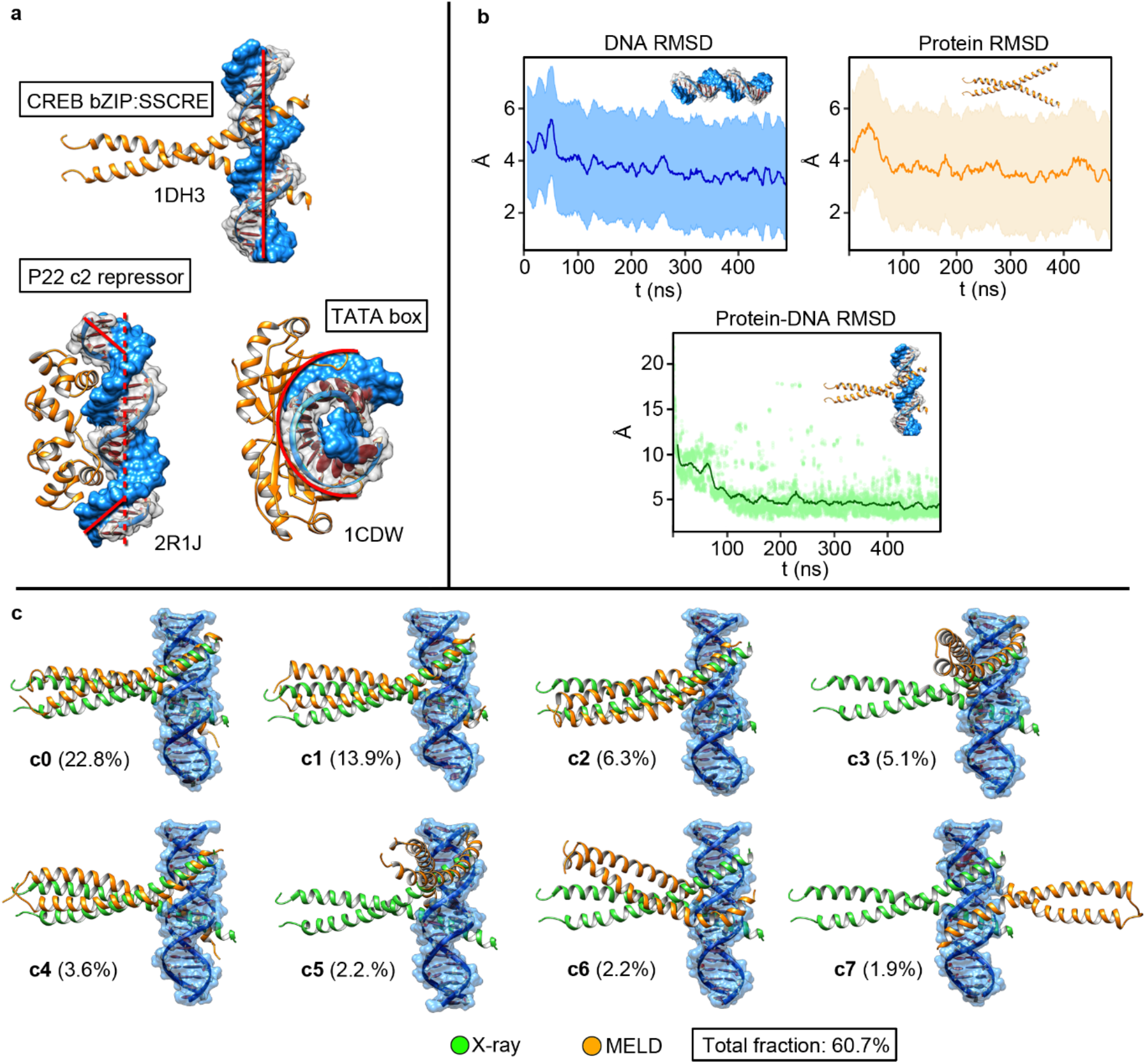
(a) Ribbon-surface representations of the three TF-DNA complexes used in this study (bZIP, P22 and TATA-box) including their PDB accession codes. (b)DNA, Protein and Protein-DNA RMSD plots. Even when using restraints on the protein and DNA, they can fluctuate significantly around the reference conformation. (c) Top binding modes for the bZIP-DNA complex using the DNA consensus sequence. The binding modes were identified through clustering (centroid of each cluster is depicted) and ranked by population (in parenthesis).

We first explored MELD-DNA’s ability to capture bound conformations using generic guiding data and allowing the DNA to deform during the process. To achieve this, we introduced the generic data as possible contacts between protein residues (mapped to C_β_) and the phosphate groups in DNA (see SI for details and Tables S2-4. The protein residues were identified from the known bound conformation, while we used repeating phosphate patterns along the DNA sequence. In this sense, the data is ambiguous and noisy as there are multiple contact interpretations for every protein residue, but only a few are present in the native state. MELD-DNA is then able to sample through combinations of possible restraints and conformations that are consistent with each other, leading to sampling multiple binding modes (see Fig. 1c for bZIP protein). For each protein system we simulated the consensus sequence and several additional sequences, running Hamiltonian and Temperature Replica Exchange Molecular Dynamics simulations (H,T-REMD) for at least a microsecond. We assessed success in two ways: i) as the ability to sample experimental conformations in the ensemble and ii) by the identification of those bound conformations based on the highest population clusters.

The case of bZIP is the most representative in terms of binding mode diversity. The data is compatible with binding through the major and minor groove as well as sliding through the sequence along the grooves (see top clusters in Fig. 1c). On the other hand, a random DNA sequence with the same guiding information produces similar clusters but changes their ranking (see Fig. S7) suggesting some sequence preferences. We further tested this idea by using five different DNA sequences and restraining each of them to 10 different B-DNA metastable states (total of 50 simulations, see Figs. S6,S8). As a result, we observed that only some sequence/structure combinations led to binding, with the consensus sequence identifying the binding site most often (see SI and Figs. S6-S9).

In the cases of P22 and TATA systems, the binding is achieved through shape complementarity, requiring the DNA to sample conformations far from canonical DNA. Protein binding induces large conformational changes in the overall DNA structure by inducing correlated conformations between base pairs, rather than severely distorting each individual base pair. Thus, the structural preferences of DNA in its free form have an impact on complex formation and are useful predictors of binding preferences. While MELD-DNA trivially recovers the binding mode when we restraint DNA conformations to the bound state, it is also successful at recovering binding modes and complex structures when DNA starts in B-DNA conformation and is allowed to deform. The guiding data is enough to bring the protein close to the DNA, which induces correlated motions in the base pairs that naturally adopt bound conformations. This approach recovered the bound conformation for all sequences tested (see Table S1), with small variations in the binding mode.

In the case of P22, binding is characterized by two interface regions along the DNA helix, where a valine in each of the binding site induces a bend in the DNA. Our simulations, guided by the ambiguous data, correctly identified this binding mode (see Fig. S10), and identified the native conformation as the highest population cluster (see Figs. S10,S11). On the other hand, the TATA box system was the most challenging one, as it introduces the largest degree of DNA deformation of all three systems. Despite this, MELD-DNA samples the experimental structure. However, we were not able to identify this conformation as the highest population cluster (see Fig. S13). Further inspection revealed that the experimental binding mode is sampled with high population in intermediate replicas, therefore the lowest replicas favor a more kinked DNA with a larger protein-DNA interface (see SI Figs. S12-S14).

For bZIP, binding simulations were enough to distinguish the consensus binding sequence from other binding poses, based on the population of the top binding mode. For TATA and P22 the nature of the data is compatible with deformation of the DNA, resulting in similar binding modes independently of the sequence.

We introduce competitive binding simulations to disambiguate binding preferences for P22 and TATA (see computational methods in SI). In this regard, we have previously shown that the MELD relative binding free energy are related to the binding free energy between the two structures^[40]^ In competitive binding, the system consists of the protein and two DNA molecules. The two DNA structures are restrained far away from each other (between 50 and 100Å from each). These restraints prevent the protein from interacting with both DNA sequences simultaneously. The guiding information is now more ambiguous: every restraint can now be satisfied between the protein and either DNA molecule (see Tables S5 and S6). At high replica indices the protein is far away from both DNA structures, while at low replica indices the protein is bound to either DNA structure. The non-bound sequence remains in its reference state (at distances greater than 50Å) sampling mostly B-DNA conformations. Analyzing the lowest replica shows the protein “jumping” from being bound to one structure or the other. A simple ratio of the bound population to each DNA structure yields binding preferences (see SI and Fig. 2). ^[40]^

**Figure 2.**
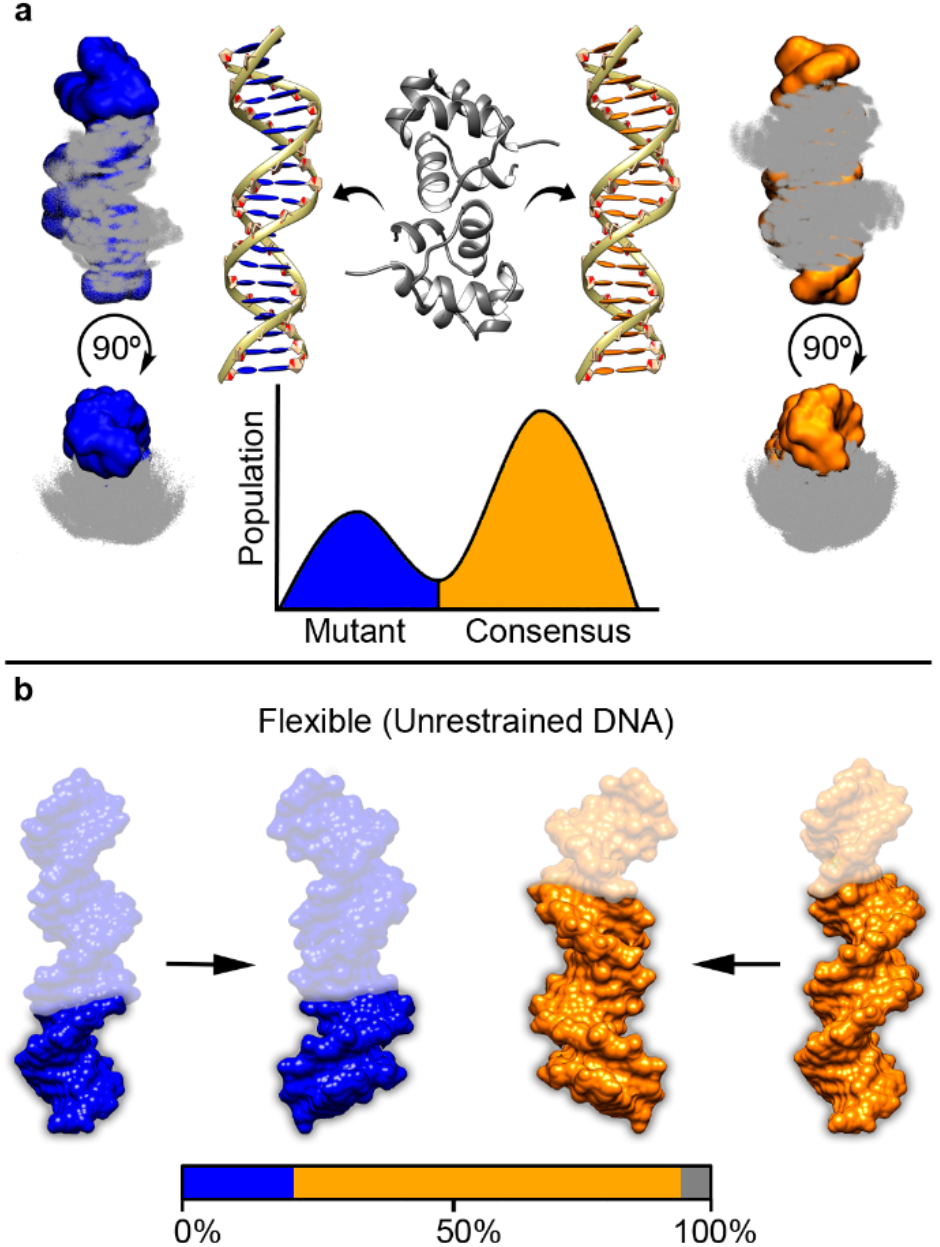
P22-DNA sequence competition. (a) MELD simulations in which the protein is directed to bind to two different DNA sequences (blue and orange). The ratio of populations where the protein is binding each sequence is related to the relative binding affinity. (b): When the DNA is allowed to freely change its conformation during competitive binding, the consensus sequence has a marked preference for binding over the mutant (population in each mode in the bar, with grey depicting non-native binding modes.

The approach is currently qualitative as it is not sensitive to large free energy differences – where only one structure might be bound at the lowest replica. The method can readily be made more quantitative by using information from all replicas with proper reweighting (e.g. using the multistate Bennett acceptance ratio), however this was out of the scope of the current work.^[41]^

We first tested simulations in which we used the consensus sequence for both DNA molecules. The expected outcome is that the protein should bind equally to each sequence and is thus a test of the expected errors as well as possible systematic errors arising from the setup conditions. The observed ratio of 57/43 populations was close to the expected value for P22 (see Fig. S12C). However, for the TATA system unbinding events were few, resulting in low number of exchanges between replicas and a systematic preference for one sequence. We suspect this is a consequence of the tighter binding discussed above and thus did not pursue this system further. We focus further discussion in the P22 system.

We extended the competitive simulations to all six sequences used in the study against the consensus sequence. In all cases the consensus sequence was heavily preferred over the other sequences (see Fig. S15C), showing the promise of the approach. Since the binding mode observed in all cases was similar, we decided to further test the importance of the shape readout mechanism. Shape readout proposes that the binding free energy is in part modulated by the ability of a DNA sequence to adopt its bound conformation. We thus carried out simulations in which the DNA was restrained to their bound conformation. We expect that in these conditions the shape readout cost is prepaid, and it should lead to closer binding populations between the consensus and mutant sequences. Figure S15B shows that this is true for some of the cases, although it is not consistent amongst all sequences. In particular, the sequence labeled ‘random’ exhibits a substantial drop in bound population. We performed further simulations for this system competing with the consensus sequence and restraining the structure of each sequence to either the holo (bound) or apo (unbound) DNA conformation (see Fig. 15A). As expected, competing the apo and holo conformations returned a massive preference for the holo conformation irrespective of the sequence. Surprisingly, when both structures are in the apo form, the preference seems to shift towards the random sequence. This would indicate that the B-DNA conformation of the random sequence has a higher initial propensity for the approach of P22, but as the protein starts interacting this sequence lacks the flexibility of the consensus sequence to bind efficiently. While this hypothesis will require further testing, and a broader benchmark set it is representative of the type of insights that MELD-DNA can provide.

Overall, the MELD-DNA methodology we presented herein fills a gap in computational tools that predict protein-DNA binding. We have shown that the method is sensitive to sequence and structural preferences and is thus a promising new approach to study this type of systems. The MELD code is freely available through github as a plugin to openmm^[42]^ The observed difficulty in promoting unbinding events for the TATA systems points at possible force field issues either in the implicit solvent or protein/DNA force field combination that warrant future investigation.^[43,44]^ Improvement in this area will contribute to more robust biophysical computational methodologies. We believe the physics-based insights MELD-DNA can provide will advance our understanding of protein-DNA interactions and our ability to simulate events related to supramolecular chemistry. Increasing our structural knowledge and sequence binding structural preferences combined with other factors that affect *in vivo* binding (e.g. chromatin state and accessibility) can bring new understanding to the molecular mechanisms that orchestrate gene regulation.

## Acknowledgements

AB and AP thank the hipergator computational resources and startup funding from the University of Florida.

