## Supplementary material for "MELD-DNA: A new tool for capturing protein-DNA binding": SI Methods and figures

**Abstract:** Herein we present MELD-DNA, a novel computational approach to address the problem of protein-DNA structure prediction. This method addresses well-known issues hampering current computational approaches to bridge the gap between structural and sequence knowledge, such as large conformational changes in DNA and highly charged electrostatic interaction during binding. MELD-DNA is able to: i) sample multiple binding modes, ii) identify the preferred binding mode from the ensembles, and iii) provide qualitative binding preferences between DNA sequences. We expect the results presented herein will have impact in the field of biophysics (through new software development), structural biology (by complementing DNA structural databases) and supramolecular chemistry (by bringing new insights into protein-DNA interactions).

**Table of Contents**

**Computational Methods**
**The MELD-DNA approach** Page 2
**Protocols for Nucleic Acid binding** Page 5
**Systems under study** Page 6
**Protein-DNA restraints** Page 6
**Competitive binding simulations** Page 8

**Results and Discussion** Page 9
**Supplementary Tables and Figures** Page 14
**References** Page 34

### Computational Methods

#### The MELD-DNA approach

*Introduction*
The larger number of degrees of freedom in Bio-macromolecules limits conformational sampling. Not all conformations are equally likely and for many applications, only the most relevant regions or metastable states in the energy surface are of interest. Molecular Dynamics (MD) based techniques are useful tools to explore metastable states while learning about their relative importance, especially with the advent of GPU enabled simulations. Even with long simulations, the study of large conformational transitions (e.g. protein binding or folding) require more advanced sampling techniques that can explore the energy surface more efficiently[1,2]

MELD-DNA builds on the successes of the MELDxMD approach[2–11] to combine general information with a molecular dynamics engine. MELD-DNA uses information to reduce the size of the conformational landscape to explore and replica exchange for more efficient sampling[12,13] When all the data is accurate and can be simultaneously satisfied, it greatly reduces the conformational landscape to focus sampling on specific states. However, in most cases there is a lack of such accurate data[7] Instead, we might rely on general knowledge that is compatible with multiple atomistic interpretations.
For example, knowing that a particular Transcription Factor (TF) binds DNA through the phosphate backbone is compatible with many possible binding sites along the sequence, and yet this knowledge already excludes all conformations in which the protein is far from the phosphate backbone. We consider this type of data ambiguous or noisy data, in the sense that one interpretation of the data leads to the correct bound conformation, while many other possible interpretations along the sequence lead to explore conformations that are not native-like[14] Just by examining the different interpretations of the data is not enough to distinguish conformations compatible with the native state. Introducing a physics models allows us to identify which of the possible interpretations of the data is the best, while at the same time narrowing the search to a manageable number of regions of interest in the conformational landscape. This is best framed as a Bayesian inference problem[15–17]

*Fundamentals of MELD-DNA*

MELD-DNA represents an extension of MELDxMD[7] which is our implementation of the Bayesian approach into a MD pipeline (Equation S1 and Figure S1).

The *prior* (p(x)) is based on an atomistic force field (the physics model) and the *likelihood* (p(D|x)) represents *the best interpretation* of the data given given the current structure. A key point here is to account for the noise in the data. Thus, rather than using all the data simultaneously to guide sampling, only a subset of the data will be enforced at any point. At each step all the data is evaluated, and the subset with greater agreement with the current structure is used until the next step, where the calculation is repeated. In this way, no information is ever discarded.

The agreement is evaluated by modeling the data as flat-bottom harmonic restraints. The flat-bottom region ensures there is no energy penalty when there is agreement between a restraint and the structure. For the subset active, deviation from the flat-bottom region will have associated forces that will help guide sampling combined (together with the force field) until the next timestep. Combining the *likelihood* and the *prior* we obtain the *posterior* distribution, p(x|D). This is the ensemble that is produced by MELD-DNA. Analysis of those ensembles using statistical mechanics produces the data subset most compatible with the physics model as well as structures that represent the most likely state (in our case the bound state).

$$p(x|D) \sim p(D|x)p(x) \qquad \text{(Equation S1)}$$

The MD engine generates new structures according to Langevin dynamics, using the parmBSC1 force field for nucleic acids[18–20] the ff14SB side force field for the protein[21,22] and the GBneck2Nu implicit solvent model[23,24] How much data is to be trusted (the size of the subset) is given by an accuracy parameter, that is chosen based on the type of data. In this work the restraints were modelled as flat-bottom harmonic restraints between pairs of atoms in the protein-DNA system, where one atom comes from the protein and one from the DNA.

*MELD-DNA data organization*

We define two degrees of hierarchical organization in the data to better account for ambiguity and noise in the data: groups and collections (see Figure S2). One or more restraints belong to a group. One or more groups belong to a collection. Groups and collections have independent accuracy parameters[7]

As a practical example, a group might contain all the possible pairings of a protein atom with all the phosphate atoms in a DNA sequence, each of those represented as a flat-bottom harmonic restraint where no energy penalty is given below a certain distance and, beyond that threshold, it increases quadratically. Finally, after a second threshold is reached, linearly. We impose the accuracy so that only one restraint is enforced in this group. Similar groups will include information about other protein-DNA interactions. Out of all the possible groups, the accuracy parameter determines the total number of protein-DNA contacts we are enforcing (the collection).

The flat-bottom region of the restraints ensures that if a subset of data is satisfied, the total energy is the same as in the original force field, leading to the exponential weight for those regions being identical as in the unbiased distribution. For regions that do not agree with any subset of the data, the restraint energy will be positive, resulting in a higher total energy than in the original force field and a lower exponential weight. Adding over all possible states results in a smaller partition function than in the original force field. The regions that agree with subsets of data will thus have a higher probability that in the original force field. Furthermore, the ratio of probabilities for regions compatible with data in this biased distribution is the same as in the unbiased force field[14]

*MELD-DNA uses H,T-REMD for enhanced sampling*

Biasing restraints increase the frustration of the energy landscape, requiring advanced sampling strategies to explore it. We use a Hamiltonian and Temperature Replica Exchange for this purpose (H,T-REMD)[13] To assign which temperature and Hamiltonian will be assigned to each replica we first map the number of replicas to a value alpha (α) that goes from 0 to 1 (so as to normalize for any simulation with different number of replicas). We then assign a function that maps an alpha value to the temperature and Hamiltonian to use in that replica (e.g. geometric scaling of temperatures). The functions used to choose temperature and Hamiltonian as well as their parameters give rise to different MELD protocols – not all of which are equally efficient. The goal for MELD-DNA protocols is to balance exploration *vs* exploitation in the system by favoring binding/unbinding events. At $\alpha=1$, the system is in full exploration, having high temperatures and no biasing restraints. At $\alpha=0$, the system is in exploitation mode, sampling at room temperature and with biases enforced strongly. At the end of the simulations, statistical mechanics treatment of the ensembles at lowest temperature produces the most likely structures compatible with the force field and data provided. Effectively, the REMD approach at high temperatures allows jumping between different regions of the conformational landscape and the low replica indexes exploit those minima. In practical terms, hierarchical clustering using a similarity measure (RMSD) and an ε of 2 is performed to identify the most likely conformations (the cluster with the highest population)[25]

#### Protocols for Nucleic Acid binding

*The GBneck2nu implicit solvent for Nucleic Acids*
Replica exchange protocols scale with the number of atoms in the system. This makes binding simulations of protein-DNA systems prohibitive as we need to accommodate solvent boxes large enough to fit unbound conformations. Recently, we developed an implicit solvent model for protein-nucleic acids[24] based on the GBneck2 approach[23] This solvent model built upon the success for protein systems allows the simultaneous simulation of protein-nucleic acids systems.

*Internal restraints keep the DNA and protein from denaturing*
Increasing the temperature in the replica exchange ladder rapidly leads to DNA and protein denaturalization. To favor binding/unbinding without denaturing these molecules we impose internal restraints. For proteins, we calculate internal $C_\alpha$-$C_\alpha$ distances and impose them through-out the trajectory to limit unfolding. This is implemented as a collection with high accuracy (95%) to allow flexibility. All $C_\alpha$-$C_\alpha$ distances below 8Å in the crystal structure are selected and enforced using flat-bottom harmonic restraints with a flat-bottom region expanding 1Å in each direction of the calculated distances.
In a similar way, we impose hydrogen bond restraints across Watson-Crick pairs to prevent the two strands from separating. This type of simulations allows the deformation of the DNA between unbound (B-DNA conformations) and bound conformations. The combination of high temperature and restraints with the DNA will sometimes lead to the exploration of non-canonical DNA conformations. In particular we observe a tendency to over-favor protein-DNA contacts, especially in long DNA sequences. These structures will be useful to parametrize future GB models. We compare these binding states with simulations in which we add flat-bottom harmonic cartesian restraints to keep DNA near a starting conformation (with a 3.5Å harmonic region for each heavy atom; $k=250$ kJ/nm$^2$). These type of cartesian restraints further allow us to answer the question of shape *vs* sequence preferences during binding. Even when using these cartesian restraints the DNA samples a wide RMSD distribution around the initial structure (see Figure 1).

#### Systems under study

We chose three protein-DNA systems according to the degree of deformation for the DNA in the bound structure: bZIP, P22 and TATA (see Figure 1a). For each system we explored the different binding modes of the consensus sequence as well as other generated sequences (see Table S1). We also performed simulations for each system in which we either let freedom to the DNA to change between B-DNA and bound conformation or restrained the DNA to either the unbound or bound conformation.

#### Protein-DNA restraints

The information is chosen to be generic in terms of allowing diverse binding modes. On the protein side: Our method presumes knowledge of the binding region, which usually can be extracted from either an experimental structure (e.g. X-ray) or binding site prediction software[26] On the DNA side: We choose the phosphate group (P, OP1, OP2) as possible interaction sites that are general to all bases. In most cases, TF-proteins bind through more than one region, thus we take advantage of this feature to remove some ambiguity in our data. Nevertheless, when binding through both regions, the distance between the binding sites needs to be respected in the DNA geometry. In the following sections we outline the specific restraints used in each protein-DNA system.

***bZip***

The protein binds as two alpha helices, each of which interacts with a different region of the DNA. We selected the $C_\beta$ residues from the termini domain of each helix (corresponding to residues 4-17 and 60-74 respectively) and a region of DNA to guide possible interactions. The first domain was set to interact with DNA residues 121 to 125 and the second domain with residues 116 to 120 (see Figure S3 and Table S2).

Each of the two domains was entered as a collection, where all possible combinatoric pairings between residues in the DNA and protein were allowed (70 restraints in the first collection and 75 restraints in the second one). We asked MELD-DNA to satisfy at least 5 groups in each collection. By choosing restraints in this way, we guide the protein to the phosphate backbone of only one DNA strand. Thus, the system is able to bind through the major or minor groove and is also able to shift a few residues up and down through the DNA. All restraints were set as flat-bottom harmonic restraints with a 6Å flat-bottom region, 6-7Å harmonic region and linear beyond. A 250 kJ/nm$^2$ force constant was used, amounting ~kBT penalty at 1Å away from the flat-bottom region.

We found that the current force field combination favored compact conformations of bZip, maximizing contacts with the protein, when we allowed full flexibility of the DNA. Thus, we chose to restraint the end-to-end distance by adding distance restraints between end-of-strand phosphates. In these systems, the starting distance between phosphate groups from different strands was around 63Å. We enforced a harmonic restraint with a flat bottom region between 55Å and 65Å. The restraint was harmonic above (/below) those values up to 67 (/53)Å and linear beyond 69 (/51) Å. We used a force constant of 250 kJ/nm$^2$ which represents about ~kBT penalty at 1Å away from the flat-bottom region.

***P22 c2 repressor***

Binding happens through two distinct domains, interacting with regions of the DNA far away in sequence. We follow the same procedure as in the case of bZIP, where now we select phosphate groups on either DNA strand and create combinatorics between several protein residues ($C_\beta$). We keep a flat-bottom region of 8Å, with the restraint penalty increasing quadratically between 8 and 9Å and linearly beyond (same force constants as with bZIP). The combinatorics are shown in Figure S4 and in Table S3. For each domain (implemented as a collection in MELD-DNA) we have six different groups, each consisting of several possible protein-DNA restraints. We ask that 3 groups be satisfied at any time and one restraint should be satisfied inside those groups. Following the MELD-DNA REMD procedure the active set of restraints changes through the trajectory.

We simulated six different sequences. The consensus sequence has three regions of interest: two binding regions and a bridge region between the two. We chose 4 sequences to change only one of those regions in each case (see Table S1) and added a random sequence for completeness. In all cases cartesian restraints were used to keep the DNA close to the bound conformation. In this protein-DNA system (PDBID: 2R1J) the binding is driven by shape complementarity, successfully identifying similar binding modes in all cases. Thus, we set up competitive binding[27] simulations (see below) to learn about the preferences during binding of P22 for sequence/conformation.

*TATA box*

The TATA system is characterized by a large surface interface between the protein and nucleic acid extending close to 10 basepairs – with nearly 30 possible $C_\beta$-P contacts. We set the accuracy so that as many as 20 contacts between $C_\beta$ and P be satisfied. We introduced ambiguity by representing contacts as distances closer than 8Å, see Table S4 and Figure S5. Simulations in which we allow the DNA to deform starting from standard B-DNA conformations were able to deform to the bound conformation in the presence of the protein and this data. However, due to the large binding interface, it was difficult to explore unbinding events. To promote them, we used an umbrella sampling potential that monitored the average distance between the DNA and the protein. While this promoted unbinding from the native site, it did so by sliding along the DNA, to maximize protein-DNA interactions, resulting in a bottleneck between replicas that where completely unbound and those that were misbound/bound.

For simulations in which we restrained the conformation to the bound state, we performed standard MD simulations for the different sequences in explicit solvent, using the conformation of the DNA at the final stage of a 500ns long simulation as the reference bound for each MELD binding simulations. While the conformation of the protein in the equilibrium trajectories remained similar (within 1.5Å to the native), the nucleic acid conformation could change significantly (up to 4Å RMSD) depending on the sequence. As before, we performed simulations for each sequence restraining each heavy atom in the DNA to within 3.5Å of their initial position using flat-bottom harmonic restraints and hydrogen bond distances between pairing nucleotides. The protein was kept from unfolding at high temperatures by initially calculating all $C_\alpha$-$C_\alpha$ contacts below 8Å and enforcing them during the simulation using flat-bottom harmonic distance restraints.

#### Competitive binding simulations

The previously described simulations allow us to identify preferred binding conformations given a sequence and data. However, it does not allow us to directly compare simulations in which the binding site is identified. This is because the replica exchange ladder does not provide a free energy of binding but rather a free energy with respect an unbound reference state, which is different for every sequence. On the other hand, competitive binding simulations[3,10] allows us to recover relative binding free energies. In these simulations the system is composed of two DNA sequences and the binding protein. The data introduced guides to binding to either DNA duplex with equal probability, requiring that when the protein is interacting with one DNA sequence, the other sequence is far away in a reference state. The reference state is chosen to be the same for each sequence in such a way that counting populations of binding to each sequence at the lowest replica provides a relative binding free energy[3,10] For the case of bZip we did not run these simulations as the initial simulations were already able to distinguish differences in the binding mode.

For P22 and TATA these binding simulations allowed one added layer of comparison. In both cases the DNA has to change conformation significantly with respect to a standard B-DNA conformation. The free energy of binding has contributions from the change in conformation in the protein, DNA and from their interaction (see Equation S2).

$$\Delta G_{bind} = \Delta G_{protein} + \Delta G_{DNA} + \Delta G_{interaction} \qquad \text{(Equation S2)}$$

The free energy contribution for the protein deformation is similar when binding different DNA sequences. However, the free energy for deforming DNA to their bound conformations changes significantly according to sequence dependent properties[28–30] The interaction free energy will depend mostly on whether the interaction takes place through specific sites present on only some sequences or through general features (e.g. DNA backbone). By restricting conformations of the DNA to either their apo/holo conformation or allowing full flexibility to the DNA the simulations can discern the role that shape/sequence recognition play in the binding process.

*P22 c2 repressor*

We compete pairs of sequences, running four experiments for each pair where the DNA is restricted to a region of conformational space (either bound or unbound) and one experiment in which the DNA is allowed to freely deform (see Table S5). For the restrained simulations we perform four experiments, including:

(i) Apo (unbound conformation) *vs* Apo sequence
(ii) Holo (bound conformation) *vs* Holo sequence
(iii) Holo *vs* Apo sequence

These simulations aim to tell us about the ability of the protein to recognize sequence or shape and the ability of a particular DNA sequence to deform to the bound conformation. For the simulations in which the DNA is free to deform, we added a flat-bottom harmonic distance restraint to keep the distance between the two DNAs from coming close with each other or too far away from each other. The blat bottom region is set up to keep the two DNA sequences at a distance between 50 and 70 Å from each other.

*TATA box*
We explored several competitive binding protocols for TATA similarly to the P22 system. However, we found this system to be more difficult due to: (1) the large interface region resulting in a lower number of transitions (longer convergence) and (2) the strong shape complementarity in the system. In this case, the data together with the protein favor a conformational change that should be difficult to explore for some of the sequences simulated. Although we mention this for completion, we believe this system will require more advanced protocols for capturing relative binding free energies.

#### Results and Discussion

**Results from Binding simulations**
For each of the three systems (see Figure 1a) we ran simulations based on the consensus sequence as well as a few additional sequences (see Table S1). We defined the noisy information for the three systems according to the methods section and applied the same information for all sequences. In all three cases the consensus sequence successfully sampled the native state. Internal restraints (see Online Methods) prevent unfolding of the protein and melting of the DNA strands while allowing them a large range of conformational flexibility (see Figure 1b). The binding/unbinding process is mapped to the replica exchange ladder, unbound states are sampled at high replica indices and bound states at the lowest replica indices. Clustering of the lowest temperature replicas allows us to identify the preferred binding conformation (see Figure 1c). Our goal is to (1) identify bound conformations and (2) identify differences in bound conformations for different DNA sequences. We find that shape complementarity drives P22 and TATA binding, whereas sequence recognition drives bZip binding. We provide details on each system below.

*CREB bZIP*
The bZIP super-family of TF is large, involving different coil-coil motifs and selectivity for different DNA sequences. For the system we study (PDBID 1DH3),[31] binding involves a coiled coil motif that binds the palindromic consensus sequence **TGACGTCA**[31] The two long helices dimerize through a leucine zipper mechanism, with a hinge point close to the binding site. Binding takes place through the major groove, in which each coil binds to one of the two recognition regions (TGAC or GTCA). The information we used to drive binding ambiguously directed the binding region of the protein to these two regions of the DNA. In the crystal structure, DNA retains its characteristic B-DNA structure in its bound conformation. Given that the binding retained a B-DNA conformation, we were interested to see if (1) we could sample and identify the correct binding mode, (2) observe differences with other sequences and (3) achieve binding of the consensus sequence restraining the conformation to different initial structures.

We simulated 5 sequences in which the consensus region was mutated to a different sequence (Table S1) in explicit solvent to extract the top two representative structures from each (a total of 10 structures). Each representative structure was the centroid of the top two clusters (labeled c0 and c1 in Figure S6). For each possible sequence we set up 10 simulations, each starting from the corresponding conformation, and restraining the structure close to the initial structure using flat-bottom harmonic restraints. This corresponds to 50 simulations, each performing 1 μs long H,T-REMD simulations with 30 replicas. Each of those figures shows the RMSD of the DNA (protein) *vs* the protein-DNA interface RMSD. Each column is restrained to the same conformation and each row keeps the same DNA sequence (see Table S1).

All plots show the results from MELD-DNA simulations using the lowest replica. Figure S6 shows that in all cases the DNA fluctuates around 1.5Å from an average position, while sampling different bound/misbound conformations. The distribution inside the grey box indicates sampling the experimental binding mode. For the consensus sequence, the experimental binding mode is sampled in high populations in 9/10 simulations, with the highest population cluster falling in this region most of the time. Other sequences have a lower preference for the canonical binding site despite starting from the same conformation and using the same data. Conformation random1-c1 merits a special mention: only using the sequence from random1 does this conformation allow sampling of the experimental binding mode. For this sequence we find a significant amount of binding through the minor groove. While for Random1 sequence, the major binding mode is through the major groove as in the experimental binding mode. Furthermore, Figure 1 and Figure S7 show the differences in the top eight binding modes identified using the consensus sequence or a random sequence.

Figure S8 contains similar information to Figure S6, except now the emphasis is on the protein conformation. This highlights a larger heterogeneity in the protein conformation, with bound-like conformations of the protein in many cases that do not correspond to native binding (high iRMSD values with low protein RMSD values). This richness in possible binding modes is further seen in a histogram distribution of all replicas in the MELD-DNA approach (see Figure S9). At high replica indices, the protein can be observed far from the binding site, as the replica index diminishes, several distinct peaks representative of different binding modes arise. At the lowest temperature, there are two major peaks representing the top clusters for the consensus sequence.

*P22*

The P22 c2 repressor protein (P22) binds DNA as a dimer. Each monomer consists of a helical bundle with a valine residue giving the specificity in the interaction[32] Two VAL33 residues (one in each binding site) induce a two point of deformation in the DNA structure characteristic of this complex (see Figure 1). Although each of them introduces a small perturbation to the local structure, the probability of sampling such a conformation for free-DNA is small. The authors of the structure (PDBID 2R1J) argued that the selectivity in this complex is due to indirect (shape) readout in which a sequence consisting of 5'-TTAA-3' at each binding site had the highest affinity[32]

Binding simulations in with cartesian restraints to keep the DNA structure close to the bound conformation reveals few possible binding modes (see Figure S10). The protein backbone remains in the same conformation throughout the different binding modes, but the side chains have more conformational freedom. In particular, the two valines inserted at the major groove of the two binding sites change their spacing significantly to accommodate binding to different base pairs along the sequence (usually one base pair up or down from the canonical binding site). Despite these subtle differences, the binding mode is overall very similar in all cases due to the shape complementarity and information guiding to the binding site. As in the case of bZIP, we looked at the histograms of all replicas to quantify the different binding modes at lower temperatures (see Figure S11). Although several binding modes are observed in the middle replicas (replicas 5 to 15), one major state is observed at the lowest temperature. This RMSD distribution encompasses small translations of the protein along the DNA major groove.

To disambiguate between sequence and structural preferences we performed competitive binding simulations, where the protein is guided to bind two possible DNA sequences. By analyzing the ensemble and counting the amount of time the protein is bound to either sequence we can approximate relative binding preference. Figure 2 and Figure S15A compare the consensus sequence with a random sequence (see Table S1). When restraining both structures towards the bound conformation, the consensus sequence is clearly preferred. However, this experiment misses the DNA conformational free energy to adopt the bound conformation. Hence, we ran different simulations keeping one of the two structures restrained in its holo or apo conformation. Not surprisingly, we find that being in the bound conformation is more important than containing the consensus sequence (shape complementarity). Interestingly, our analysis seems to indicate that if both sequences are in their B-DNA conformation, the protein has some preference for the random sequence. This reinforces the view that it is not the sequence that is specifically being recognized through specific contacts (sequence readout), but the ability of a certain sequence to adopt a bound conformation (shape readout) which allows for better interactions through shape complementarity. We finally started both sequences from their B-DNA conformation and removed the cartesian restraints on the DNA. During binding the structures are able to deform to capture the correct bound state. Again, showing that the consensus sequence is preferred.

*TATA*

This protein DNA interaction[33] involves a high degree of flexibility in both the DNA and the protein. The protein binding interface is formed by β-strands tracking the DNA minor groove, flanked by two loop regions on either end that effectively wrap around the DNA minor groove. The distance between the tips of the loops can change significantly, with close distances favoring increased kinking in the DNA. Kinking reduces the stacking interactions that stabilize DNA and are therefore favored in sequences where that stacking interaction is smaller, such as the TATA sequence. The TATA binding protein (TBP) sequence is made up to two similar sequence fragments that fold into very similar structural components, each of which binds a different DNA region[33]

The information that we used for this system tracks the protein-DNA interaction along the minor groove, with enough wiggle room that the interface is not completely defined. Surprisingly, starting simulations from the B-DNA conformation and allowing flexibility on the DNA (see Online Methods) and protein allowed us to sample native conformations, with some conformations having a protein-DNA RMSD below 2.5Å (see Figure S12). However, these native-like conformations were not identified as the highest population cluster at the lowest replica-index. Instead, they were sampled through replicas 11-20, which have a higher temperature for the system. As the temperature was reduced in replicas 1-10, a different conformation emerged as the most stable under the current force field conditions (see Figure S13). The stable conformations had a protein-DNA RMSD between 3.5Å and 4.5Å, with internal DNA conformations greater than 5Å. The protein loop-to-loop distance was reduced from 30.8Å in the experimental structure to 19.0Å in our most populated cluster. This in turn increases DNA kinking (see Figure S14).

We tested the ability to sample and identify correct binding poses as well as identify sequence preferences by restraining DNA conformations close to the bound conformation (see methods). The consensus DNA sequence contains a TATAAAA sequence at the binding interface, which maps to an interaction with each structural repeat in the protein. We designed three additional changes by changing the interface in each region or both simultaneously (see Table S1). We performed binding simulations with the same protocols as with the full flexibility. In these conditions, binding of the consensus sequence maintained a loop-to-loop distance close to the experimental one (see Figure S14) and the protein-DNA RMSD for the top clusters was below 2.5Å. All four simulations identify the bound conformation, with protein-DNA RMSDs close to 2.5Å as the most populated cluster at low population, and the internal conformation of DNA deviating between 2 and 3Å from the starting conformation. However, when we look at the relative position of the DNA in the complex (see Figure S12 top panel) we see that sequences which maintain the TATA recognition site are interacting as in the experimental structures. The internal structure of the protein (Figure S12 bottom panel) shows a marked difference when it is binding DNA that is restrained to the holo-DNA structure, showing a marked lower RMSD than when the DNA is able to freely deform (blue data).

Competitive binding simulations for TATA protein did not provide further insights (see table S6). When using the bound conformation, shape complementarity and electrostatic interactions are dominating the binding. The limitations in binding free DNA get further accentuated in competitive binding, limiting the usefulness of the approach for such large deviations from the B-DNA structure with our current implicit solvent model.

Overall, TATA binding shows a behavior similarly to P22, where the binding is driven by shape complementarity. In this case, the DNA exhibits a much larger degree of deformation, giving rise to a very well-defined binding partner that is identified early in the replica exchange ladder. In a parallel way to the P22 case, removing the cartesian restraints to the DNA allows sampling of the deformation from B-DNA to the bound conformation, given the information we provide (see Figure S14). Contrary to the case of P22, competitive binding simulations are not able to identify the consensus sequence as the best binder. We believe this to be a limitation of the combination of the force field with implicit solvent, leading to improved protein-DNA interactions through shape complementarity to overshadow sequence dependent properties.

**Discussion**

Combining structural and sequence knowledge yields more accurate predictions of TF binding preferences, relevant to gene regulation[30,34–38] This sequence knowledge has been derived based on average free-DNA properties – thus structural insights help predict *where* a TF will bind, but not *how.* As a community, we lack tools to predict structural binding. The Protein Data Bank (PDB)[39] and Nucleic Acid Database (NDB)[40] provide the experimental structures for protein-DNA complexes, but this is not systematic across many sequences for a particular TF or even possible for all TF. TF bind promiscuously, allowing many protein-DNA complexes in principle, but the affinity (relative binding to sequences) can be several orders of magnitude[41] Our approach provides a tool for exploring protein-DNA binding interactions at a structural level. The tool overcomes some of the previous issues in modeling DNA flexibility[42–45] and large deformations during binding through the use of H,T-REMD. It also addresses the challenges observed in docking scoring functions for highly charged systems and successfully identifies the correct bound state for the systems under study. However, the implicit solvent used remains problematic to study some cases of extreme deformation (e.g. TATA system). Future endeavors using more accurate physical models (e.g. explicit solvent) can improve current limitations while keeping retaining the current methodological approach.

As in other MD based approaches, the success of the methodology hinges on the ability to sample multiple binding/unbinding events. In the study of protein-protein and protein-peptide systems, increasing the temperature at high replica index is enough to unbind the two molecules. For protein-DNA systems, temperature alone is not enough, limiting sampling to regions close to the first site of interaction. For some of the systems we included restraints to pull the protein away from the DNA – often resulting in bottlenecks in the REMD exchange probabilities which had to be addressed. Despite the success of the current methodology, further work is needed to optimize and increase transferability of the approach.

The choice of implicit solvent is a balance between the ability to sample conformations efficiently and using REMD – which would be computationally prohibitive in explicit solvent. In the development of the implicit solvent, parameters are optimized against a dataset of good structures and bad structures[24]. During this study we have observed that using advanced sampling (even with restraints) we sample some structures at high temperature that are unreasonable. Such structures will be useful in further parameter improvements for the implicit solvent. This might explain the lack of selectivity we observe for TATA binding, where the bound structure adopts extreme kinks.
A critical step in the work is the determinization of the data to use. For all cases we assumed the known binding region in the protein is known. We then used combinatoric rules to impose generic possible interactions along the DNA sequence through the phosphate backbone. New studies show that the binding regions in the protein can be accurately estimated for cases where no structural information is known, increasing the versatility of the current MELD-DNA approach.

#### Supplementary Tables and Figures

**Table S1.** DNA sequences used for bZIP, TATA and P22 complexes. DNA bases highlighted in orange represent the changes with respect to the consensus sequence.

| Transcription Factor | Sequence | |
|---|---|---|
| **bZIP** | Consensus | CCTTGG**CTGACGTCAG**CCAAG |
| | noCG | CCTTGGCTGAATTCAGCCAAG |
| | Random 1 | CCTTGGATGCTACGATCCAAG |
| | Random 2 | CCTTGGCGTAGCTCGGCCAAG |
| | Random 3 | CCTTGGTCTATCGGTTCCAAG |
| **TATA box** | Consensus | CTGC**TATAAAA**GGCTG |
| | Domain 1 | CTGCCGCGAAAGGCTG |
| | Domain 2 | CTGCTATAGGGGGCTG |
| | Random 1 | CTGCCGCGGGGGGCTG |
| **P22 c2 repressor** | Consensus | CATTT**AAG**ATAT**CTT**AAATA |
| | Domain 1 | CATTTCCTATATCTTAAATA |
| | Domain 2 | CATTTAAGATATGGCAAATA |
| | Bridge 1 | CATTTAAGCGCGCTTAAATA |
| | Bridge 2 | CATTTAAGCCGACTTAAATA |
| | Random 1 | CGCCATTTAGGGACGATCCA |

**Table S2**. Information used for bZIP-DNA binding. We asked MELD-DNA to satisfy at least 5 groups in each collection. Groups where defined based on all the combinatorics of residues in DNA pairing with those in the protein. Each group had 3 possible restraints between the $C_\beta$ in the protein and 3 atoms in DNA.

| Collection 1 (Domain 1) | | | | | |
|---|---|---|---|---|---|
| | **DNA-resid** | **DNA atom** | **Prot-resid** | **Protein atom** | **Distance (Å)** |
| **Groups (70)** | 121 to 125 | P, OP1, OP2 | 4 to 17 | $C_\beta$ | 8.0 |
| **Collection 2 (Domain 2)** | | | | | |
| | **DNA-resid** | **DNA atom** | **Prot-resid** | **Protein atom** | **Distance (Å)** |
| **Groups (75)** | 116 to 120 | P, OP1, OP2 | 60 to 74 | $C_\beta$ | 8.0 |

**Table S3.** Information used for the P22-DNA binding. Each group was evaluated best on one restraint and each collection was enforcing three groups according to the MELD-DNA strategy. Concretely, DNA Residue ID (DNA-resid), DNA atom, Protein Residue ID (Prot-Resid), Protein atom and distances (in Å) are indicated.

| Collection 1 (Domain 1) | | | | | |
|---|---|---|---|---|---|
| | **DNA-resid** | **DNA atom** | **Prot-resid** | **Protein atom** | **Distance (Å)** |
| **Group 1** | 21 | O5' | 49 | $C_\beta$ | 8.0 |
| | 21 | O5' | 59 | $C_\beta$ | 8.0 |
| | 21 | O5' | 70 | $C_\beta$ | 8.0 |
| | 21 | O5' | 74 | $C_\beta$ | 8.0 |

| | | | | | |
|---|---|---|---|---|---|
| | 21 | O5' | 75 | $C_\beta$ | 8.0 |
| **Group 2** | 22 | P | 49 | $C_\beta$ | 8.0 |
| | 22 | P | 59 | $C_\beta$ | 8.0 |
| | 22 | P | 70 | $C_\beta$ | 8.0 |
| | 22 | P | 74 | $C_\beta$ | 8.0 |
| | 22 | P | 75 | $C_\beta$ | 8.0 |
| **Group 3** | 23 | P | 49 | $C_\beta$ | 8.0 |
| | 23 | P | 59 | $C_\beta$ | 8.0 |
| | 23 | P | 70 | $C_\beta$ | 8.0 |
| | 23 | P | 74 | $C_\beta$ | 8.0 |
| | 23 | P | 75 | $C_\beta$ | 8.0 |
| **Group 4** | 24 | P | 49 | $C_\beta$ | 8.0 |
| | 24 | P | 59 | $C_\beta$ | 8.0 |
| | 24 | P | 70 | $C_\beta$ | 8.0 |
| | 24 | P | 74 | $C_\beta$ | 8.0 |
| | 24 | P | 75 | $C_\beta$ | 8.0 |
| **Group 5** | 25 | P | 49 | $C_\beta$ | 8.0 |
| | 25 | P | 59 | $C_\beta$ | 8.0 |
| | 25 | P | 70 | $C_\beta$ | 8.0 |
| | 25 | P | 74 | $C_\beta$ | 8.0 |
| | 25 | P | 75 | $C_\beta$ | 8.0 |
| **Group 6** | 12 | P | 69 | $C_\beta$ | 8.0 |
| | 13 | P | 69 | $C_\beta$ | 8.0 |
| | 14 | P | 69 | $C_\beta$ | 8.0 |
| | 15 | P | 69 | $C_\beta$ | 8.0 |
| | 16 | P | 69 | $C_\beta$ | 8.0 |
| **Collection 2 (domain 2)** | | | | | |
| | **DNA-resid** | **DNA atom** | **Prot-resid** | **Protein atom** | **Distance** |
| **Group 1** | 3 | P | 115 | $C_\beta$ | 8.0 |
| | 3 | P | 125 | $C_\beta$ | 8.0 |
| | 3 | P | 140 | $C_\beta$ | 8.0 |
| | 3 | P | 141 | $C_\beta$ | 8.0 |
| **Group 2** | 4 | P | 115 | $C_\beta$ | 8.0 |
| | 4 | P | 125 | $C_\beta$ | 8.0 |
| | 4 | P | 140 | $C_\beta$ | 8.0 |
| | 4 | P | 141 | $C_\beta$ | 8.0 |

| | | | | | |
|---|---|---|---|---|---|
| **Group 3** | 2 | P | 115 | $C_\beta$ | 8.0 |
| | 2 | P | 125 | $C_\beta$ | 8.0 |
| | 2 | P | 140 | $C_\beta$ | 8.0 |
| | 2 | P | 141 | $C_\beta$ | 8.0 |
| **Group 4** | 5 | P | 115 | $C_\beta$ | 8.0 |
| | 5 | P | 125 | $C_\beta$ | 8.0 |
| | 5 | P | 140 | $C_\beta$ | 8.0 |
| | 5 | P | 141 | $C_\beta$ | 8.0 |
| **Group 5** | 6 | P | 115 | $C_\beta$ | 8.0 |
| | 6 | P | 125 | $C_\beta$ | 8.0 |
| | 6 | P | 140 | $C_\beta$ | 8.0 |
| | 6 | P | 141 | $C_\beta$ | 8.0 |
| **Group 7** | 32 | P | 135 | $C_\beta$ | 8.0 |
| | 33 | P | 135 | $C_\beta$ | 8.0 |
| | 34 | P | 135 | $C_\beta$ | 8.0 |
| | 35 | P | 135 | $C_\beta$ | 8.0 |
| | 36 | P | 135 | $C_\beta$ | 8.0 |

**Table S4.** Information used for the TATA-DNA binding. A collection with a single group, in which 20 of the restraints were enforced at any step of the simulation. Concretely, DNA Residue ID (DNA-resid), DNA atom, Protein Residue ID (Prot-Resid), Protein atom and distances (in Å) are indicated.

| DNA-resid | DNA atom | Prot-resid | Protein atom | Distance (Å) |
|---|---|---|---|---|
| 6 | P | 161 | $C_\beta$ | 8.0 |
| 6 | P | 162 | $C_\beta$ | 8.0 |
| 8 | P | 175 | $C_\beta$ | 8.0 |
| 10 | P | 130 | $C_\beta$ | 8.0 |
| 11 | P | 45 | $C_\beta$ | 8.0 |
| 12 | P | 88 | $C_\beta$ | 8.0 |
| 12 | P | 90 | $C_\beta$ | 8.0 |
| 12 | P | 92 | $C_\beta$ | 8.0 |
| 13 | P | 72 | $C_\beta$ | 8.0 |
| 13 | P | 88 | $C_\beta$ | 8.0 |
| 13 | P | 89 | $C_\beta$ | 8.0 |
| 14 | P | 68 | $C_\beta$ | 8.0 |
| 14 | P | 72 | $C_\beta$ | 8.0 |
| 22 | P | 70 | $C_\beta$ | 8.0 |
| 23 | P | 71 | $C_\beta$ | 8.0 |
| 23 | P | 75 | $C_\beta$ | 8.0 |
| 24 | P | 84 | $C_\beta$ | 8.0 |
| 24 | P | 96 | $C_\beta$ | 8.0 |
| 25 | P | 41 | $C_\beta$ | 8.0 |
| 26 | P | 40 | $C_\beta$ | 8.0 |
| 27 | P | 135 | $C_\beta$ | 8.0 |
| 27 | P | 183 | $C_\beta$ | 8.0 |
| 28 | P | 179 | $C_\beta$ | 8.0 |
| 28 | P | 181 | $C_\beta$ | 8.0 |
| 28 | P | 183 | $C_\beta$ | 8.0 |
| 29 | P | 163 | $C_\beta$ | 8.0 |
| 29 | P | 179 | $C_\beta$ | 8.0 |
| 29 | P | 180 | $C_\beta$ | 8.0 |
| 29 | P | 181 | $C_\beta$ | 8.0 |

**Table S5.** Competitive binding simulation information for the P22 system. Each group contains information that can be satisfied with respect to one DNA sequence or the other. 12 of the 19 groups were enforced at any time in the simulation. Concretely, DNA Residue ID (DNA-resid), DNA atom, Protein Residue ID (Prot-Resid), Protein atom and distances (in Å) are indicated.

| | DNA-resid | DNA atom | Prot-resid | Protein atom | Distance (Å) |
|---|---|---|---|---|---|
| **Group 1** | 3 | P | 164 | $C_\beta$ | 6.0 |
| | 43 | P | 164 | $C_\beta$ | 6.0 |
| **Group 2** | 3 | P | 165 | $C_\beta$ | 6.0 |
| | 43 | P | 165 | $C_\beta$ | 6.0 |
| **Group 3** | 4 | P | 180 | $C_\beta$ | 6.0 |
| | 44 | P | 180 | $C_\beta$ | 6.0 |
| **Group 4** | 34 | P | 174 | $C_\beta$ | 6.0 |
| | 74 | P | 174 | $C_\beta$ | 6.0 |
| **Group 5** | 34 | P | 177 | $C_\beta$ | 6.0 |
| | 74 | P | 177 | $C_\beta$ | 6.0 |
| **Group 6** | 34 | P | 178 | $C_\beta$ | 6.0 |
| | 74 | P | 178 | $C_\beta$ | 6.0 |
| **Group 7** | 33 | P | 190 | $C_\beta$ | 6.0 |
| | 73 | P | 190 | $C_\beta$ | 6.0 |
| **Group 8** | 33 | P | 189 | $C_\beta$ | 6.0 |
| | 73 | P | 189 | $C_\beta$ | 6.0 |
| **Group 9** | 33 | P | 193 | $C_\beta$ | 6.0 |
| | 73 | P | 193 | $C_\beta$ | 6.0 |
| **Group 10** | 32 | P | 187 | $C_\beta$ | 6.0 |
| | 72 | P | 187 | $C_\beta$ | 6.0 |
| **Group 11** | 32 | P | 188 | $C_\beta$ | 6.0 |
| | 72 | P | 188 | $C_\beta$ | 6.0 |
| **Group 12** | 13 | P | 124 | $C_\beta$ | 6.0 |
| | 53 | P | 124 | $C_\beta$ | 6.0 |
| **Group 13** | 13 | P | 127 | $C_\beta$ | 6.0 |
| | 53 | P | 127 | $C_\beta$ | 6.0 |

| | | | | | |
|---|---|---|---|---|---|
| **Group 14** | 13 | P | 122 | $C_\beta$ | 6.0 |
| | 53 | P | 122 | $C_\beta$ | 6.0 |
| **Group 15** | 14 | P | 109 | $C_\beta$ | 6.0 |
| | 54 | P | 109 | $C_\beta$ | 6.0 |
| **Group 16** | 14 | P | 108 | $C_\beta$ | 6.0 |
| | 54 | P | 108 | $C_\beta$ | 6.0 |
| **Group 17** | 14 | P | 112 | $C_\beta$ | 6.0 |
| | 54 | P | 112 | $C_\beta$ | 6.0 |
| **Group 18** | 23 | P | 98 | $C_\beta$ | 6.0 |
| | 63 | P | 98 | $C_\beta$ | 6.0 |
| **Group 19** | 24 | P | 114 | $C_\beta$ | 6.0 |
| | 64 | P | 114 | $C_\beta$ | 6.0 |

**Table S6.** Competitive binding simulation information for the TATA binding protein. Each group contains information that can be satisfied with respect to one DNA sequence or the other. 6 of the 8 groups were enforced at any point in the simulation. Concretely, DNA Residue ID (DNA-resid), DNA atom, Protein Residue ID (Prot-Resid), Protein atom and distances (in Å) are indicated.

| | DNA-resid | DNA atom | Prot-resid | Protein Atom | Distance (Å) |
|---|---|---|---|---|---|
| **Group 1** | 28 | OP1 | 213 | OG | 5.0 |
| | 60 | OP1 | 213 | OG | 5.0 |
| **Group 2** | 23 | OP1 | 102 | NH2 | 5.0 |
| | 23 | OP1 | 102 | NH1 | 5.0 |
| | 55 | OP1 | 102 | NH2 | 5.0 |
| | 55 | OP1 | 102 | NH1 | 5.0 |
| **Group 3** | 24 | OP1 | 109 | NH2 | 5.0 |
| | 24 | OP1 | 109 | NE | 5.0 |
| | 56 | OP1 | 109 | NH2 | 5.0 |
| | 56 | OP1 | 109 | NE | 5.0 |
| **Group 4** | 24 | OP1 | 116 | OG1 | 5.0 |
| | 56 | OP1 | 116 | OG1 | 5.0 |
| **Group 5** | 10 | OP1 | 222 | NZ | 5.0 |
| | 42 | OP1 | 222 | NZ | 5.0 |
| **Group 6** | 9 | O3' | 222 | NZ | 5.0 |
| | 41 | O3' | 222 | NZ | 5.0 |
| **Group 7** | 12 | OP1 | 124 | NZ | 5.0 |
| | 44 | OP1 | 124 | NZ | 5.0 |
| **Group 8** | 8 | OP1 | 200 | NH1 | 5.0 |
| | 40 | OP1 | 200 | NH1 | 5.0 |

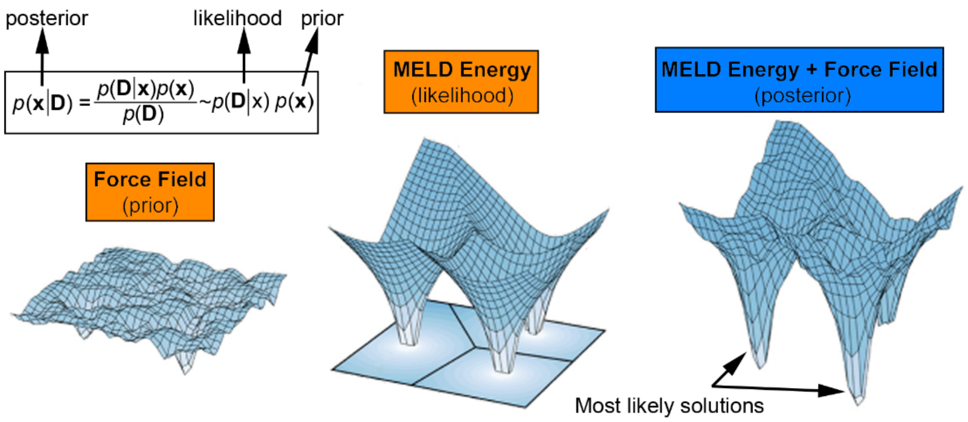

**Figure S1.** Schematic representation of MELD-DNA fundamentals. Adapted from reference[7].

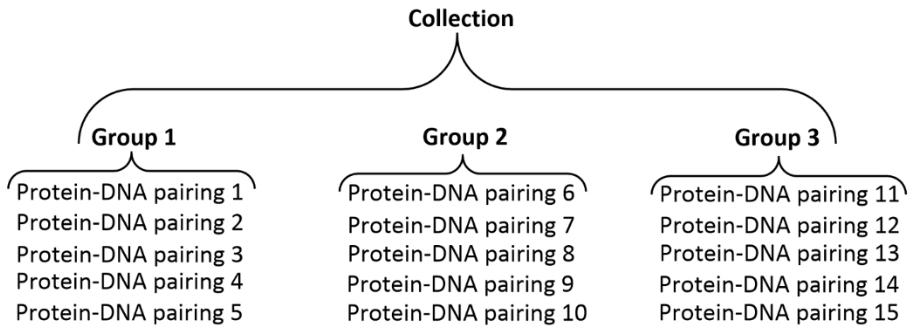

**Figure S2.** Hierarchical organization of MELD-DNA data.

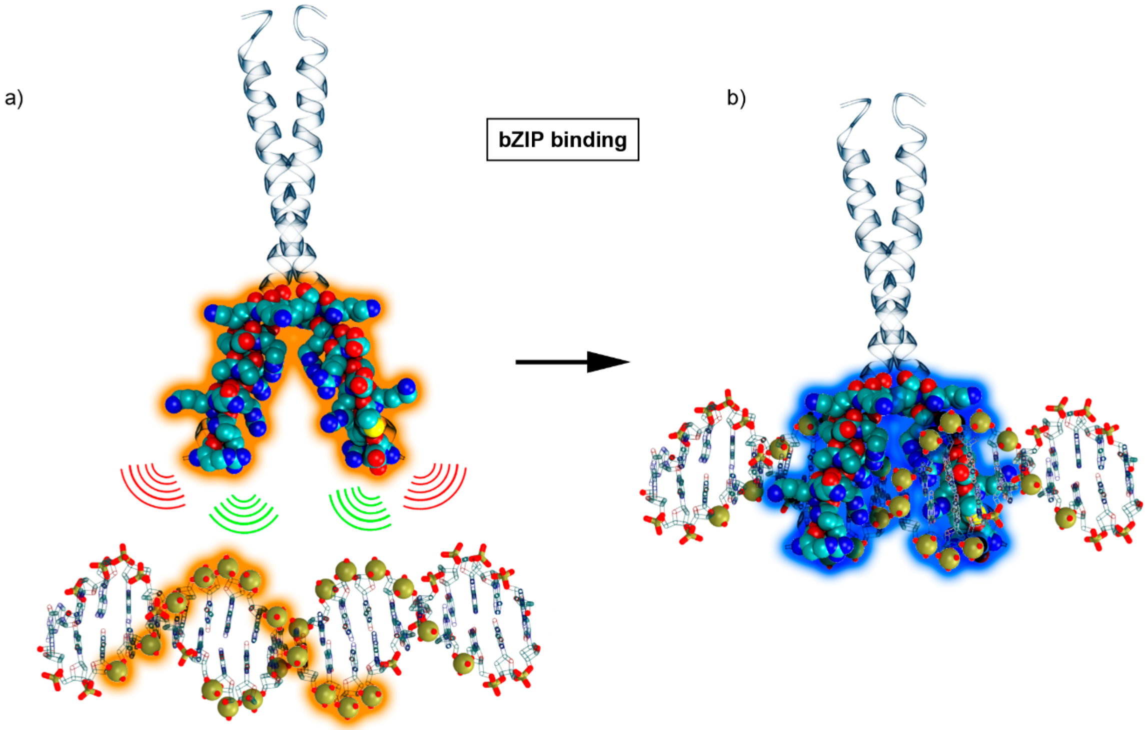

**Figure S3.** Artistic representation of ambiguity for bZIP binding. (a) Phosphate sites along the DNA sequence are combinatorically paired with $C_\beta$ atoms in the binding site of the bZIP protein (highlighted in orange). (b) The native interactions found in the experimental structure are highlighted with a blue halo.

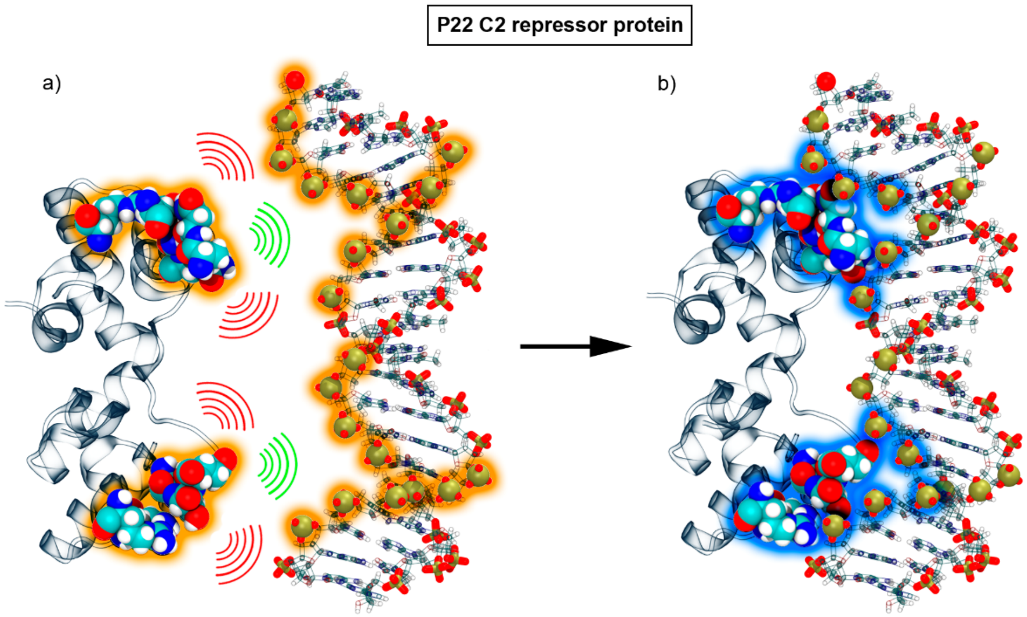

**Figure S4.** Artistic representation of ambiguity in protein-DNA systems. (a) Phosphate sites along the DNA sequence are combinatorically paired with $C_\beta$ atoms in the binding site of the P22 c2 repressor protein and highlighted in orange. The blue halo (in b) shows the native interactions undertaken between both moieties.

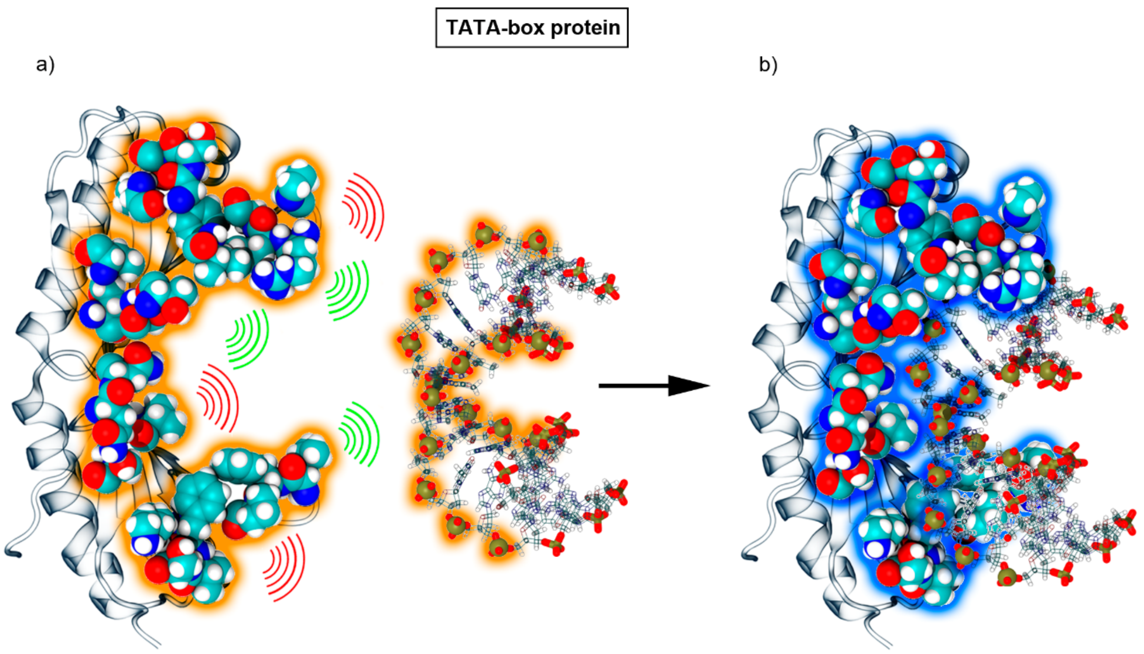

**Figure S5.** Artistic representation of ambiguity in protein-DNA systems. (a) Phosphate sites along the DNA sequence are combinatorically paired with $C_\beta$ atoms in the binding site of the TATA-box binding protein (highlighted in orange). The blue halo (b) shows the native interactions established between both moieties.

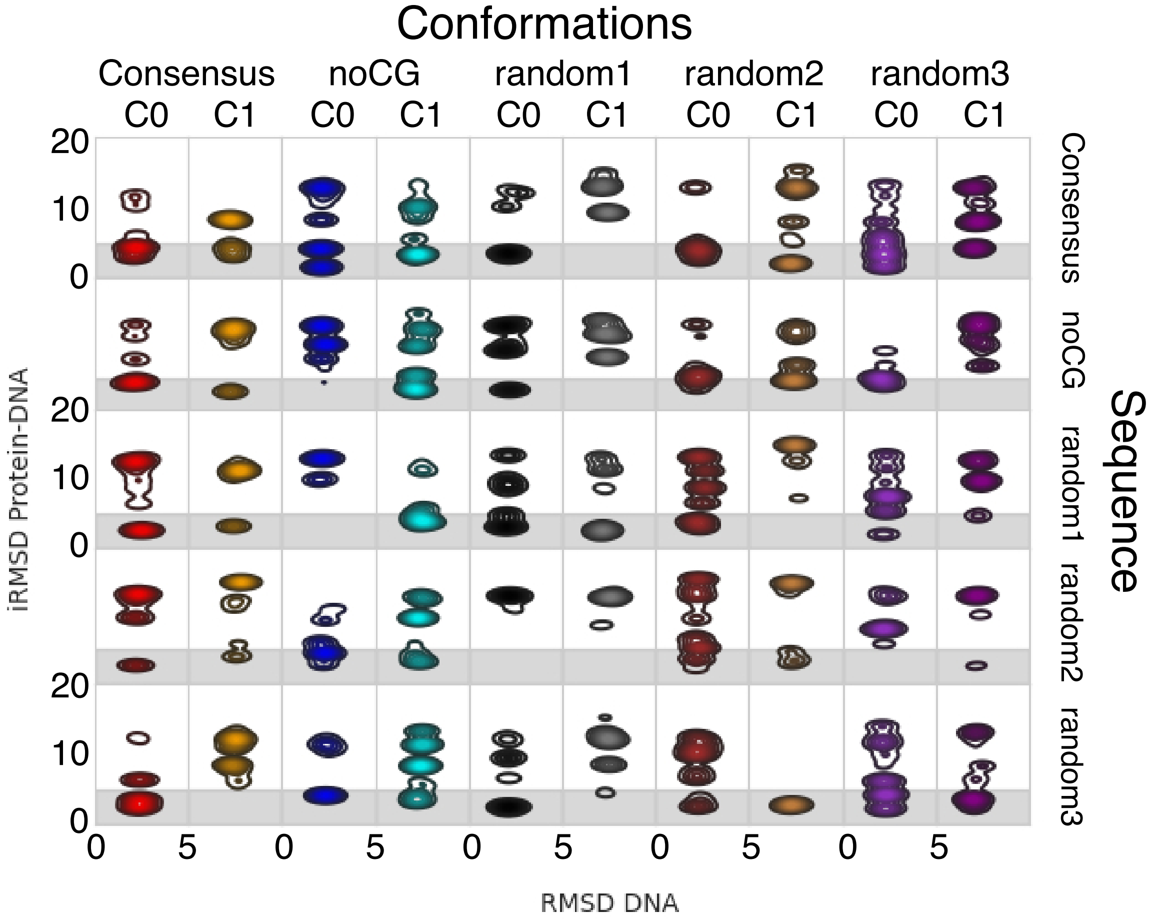

**Figure S6.** Multiple sequences and conformations are viable for Bzip-DNA binding. Each plot represents a particular sequence (denoted by the row, see labels on the right) and DNA conformation (denoted by column, belonging to clusters from free DNA simulations on each sequence). Each plot represents the interface RMSD vs the DNA RMSD. All DNA sample conformations around the initial structure, even when BZip is bound. The grey box denotes regions that we classify as successfully bound conformations, outside the grey are represents misbound conformations. All sequence/structure combinations sample multiple binding conformations. The consensus sequence is the most successful at binding in high populations in the correct binding site – only in one case the correct bound. Conformation is not sampled. Random2 sequence is the least likely to make bound conformations no matter the initial conformation – even when the correct binding conformation is sampled, it has low population.

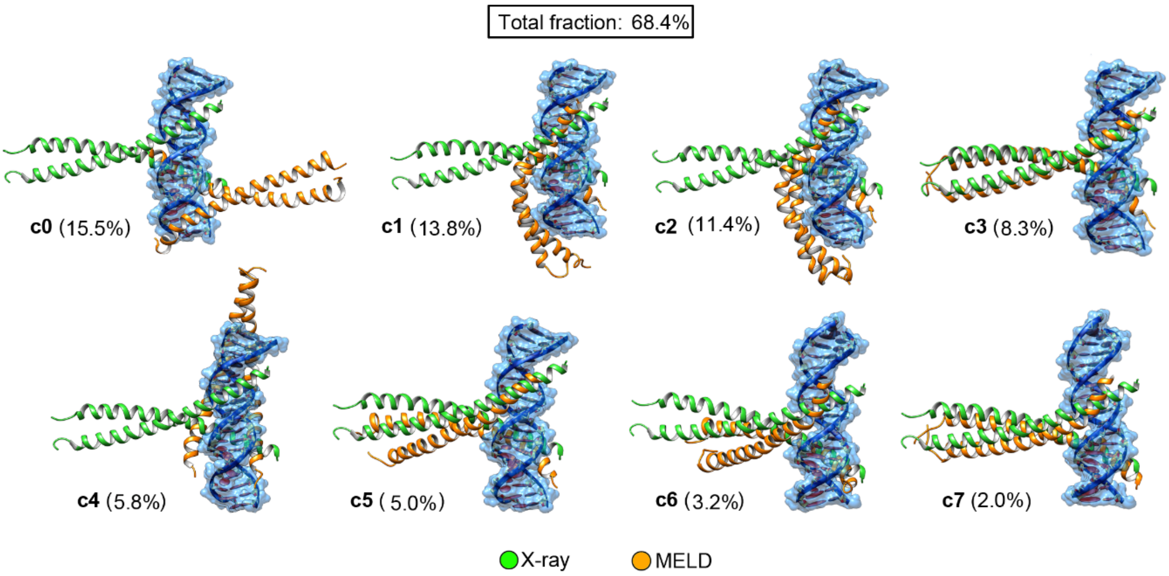

**Figure S7.** Cluster population analysis and representative structures of a bZIP-DNA complex using a random DNA sequence.

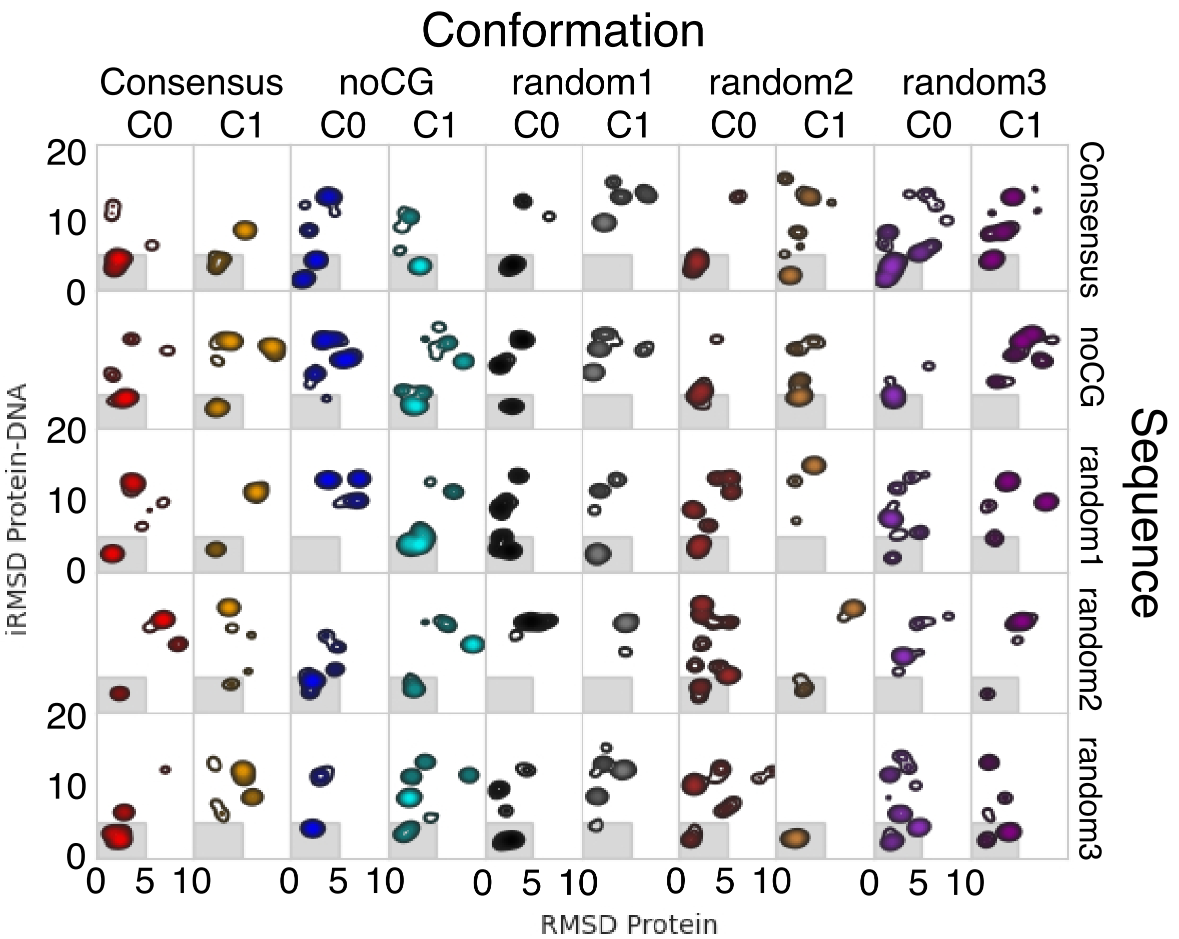

**Figure S8.** Multiple sequences and conformations are viable for Bzip-DNA binding. Each plot represents a particular sequence (denoted by the row, see labels on the right) and DNA conformation (denoted by column, belonging to clusters from free DNA simulations on each sequence). Each plot represents the interface RMSD vs the protein RMSD. MELD-DNA allows conformational freedom to the protein, which can sample a diverse set of conformations, but is mostly found in its native-like conformation when binding -- either at the correct binding site (grey box area) or others, denoted by protein RMSD below 5Å.

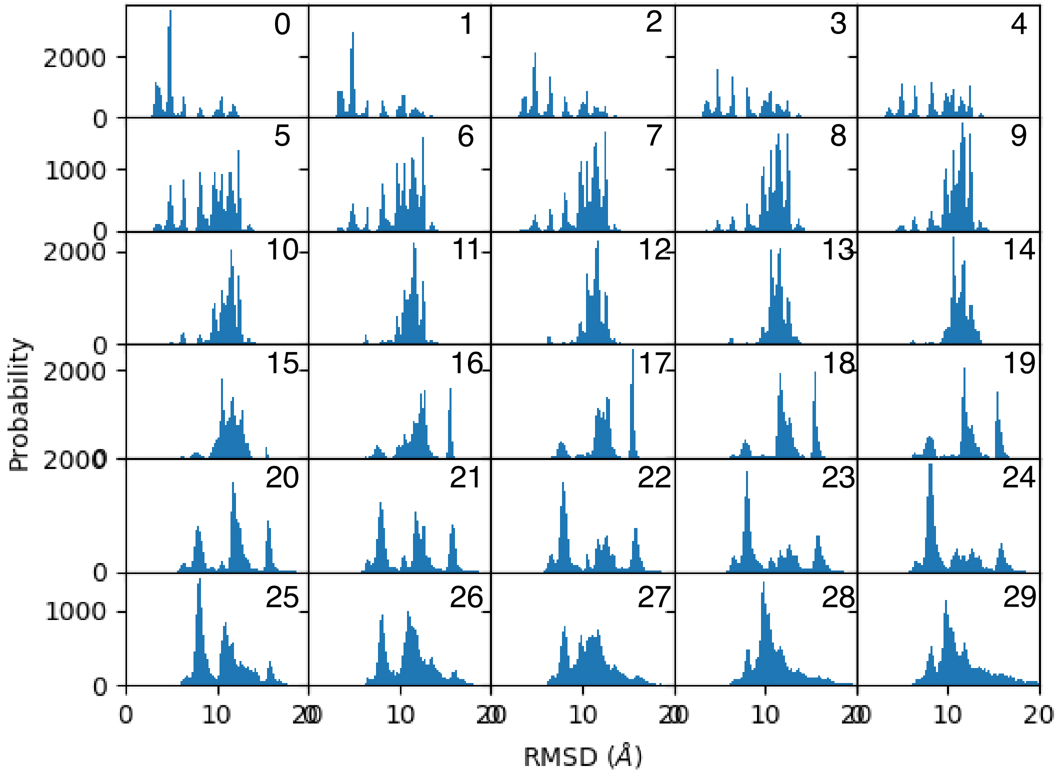

**Figure S9.** RMSD distribution for each replica in the Replica Exchange ladder for BZip. Replica 0 represents the lowest temperature and restraints enforced at the full force constant. Replica 29 corresponds to sampling at thighest temperature, with no protein-DNA restraints active. For higher replica index misbound and unbound states are sampled. As the replica index goes down, several bound and misbound states arise, with the native bound state becoming the predominant one at the lowest replica. The current plot represents the consensus sequence restraining the DNA structure to sample around the top cluster of the consensus free DNA.

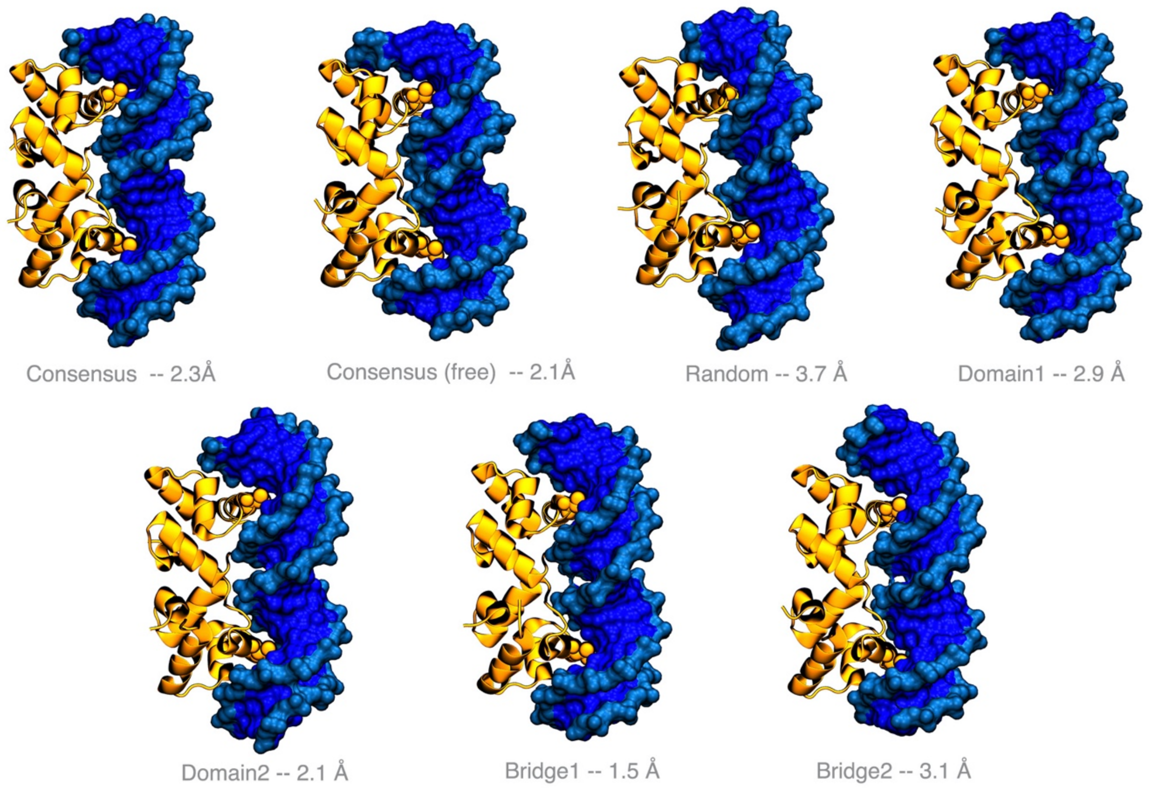

**Figure S10.** Binding modes for the different simulations in the P22 system. The RMSD is calculated for all Cα and P atoms. The structures correspond to the centroid of the highest population cluster.

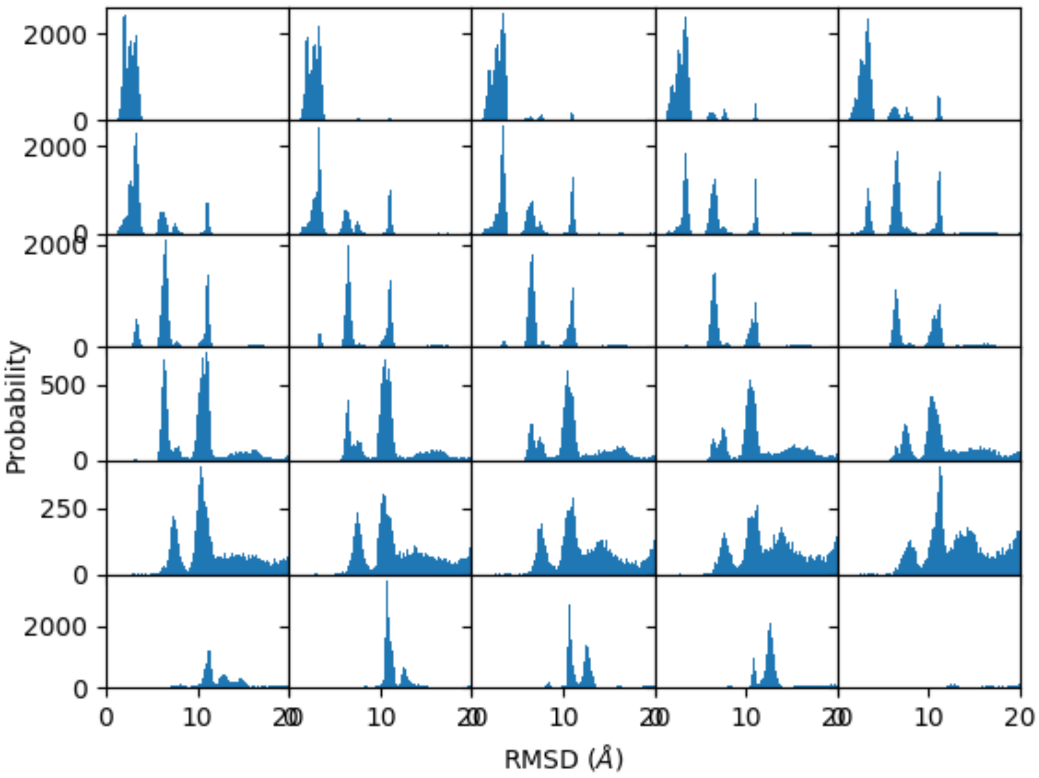

**Figure S11**. RMSD distribution for each replica in the Replica Exchange ladder for P22, following the same representation as in Figure S4. At low temperatures binding locks the structure close to the experimental conformation.

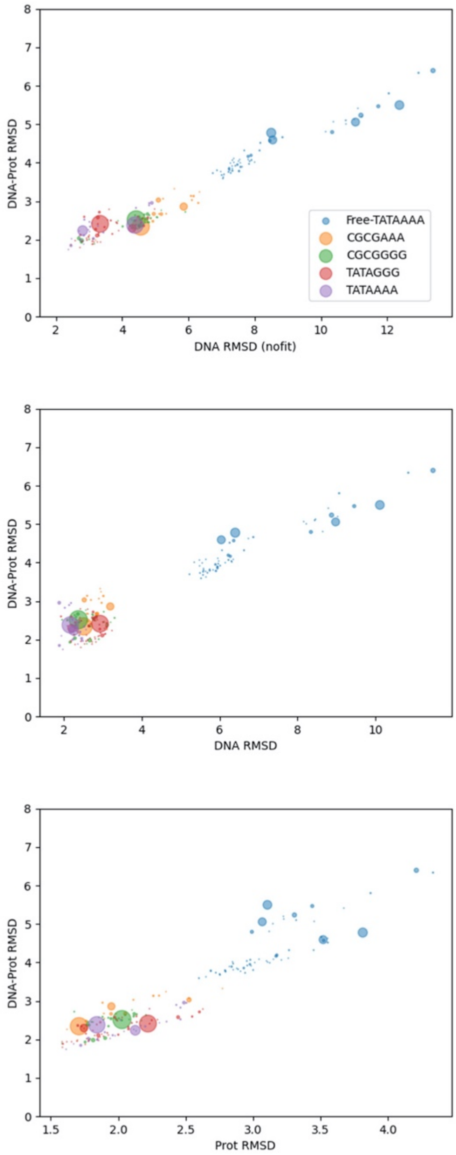

**Figure S12.** Accuracy of the top cluster in TATA-box binding simulations. Each simulation keeps the DNA restrained to the bound conformation except for the consensus sequence which is simulated restrained (purple) and starting from B-DNA (blue). Each cluster is represented as a dot, with population proportional to the size of the dot. When the DNA is able to deform, the protein wraps around the DNA major Groove more tightly, resulting in larger protein and DNA RMSD with respect to the restrained simulations. Restraining the. DNA also keeps the protein closer to its native bound conformation. When considering the relative position of the DNA to the TATA box (top panel), the consensus sequence (purple) and the mutant that conserves the TATA sequence (red) are the closest to the native binding mode.

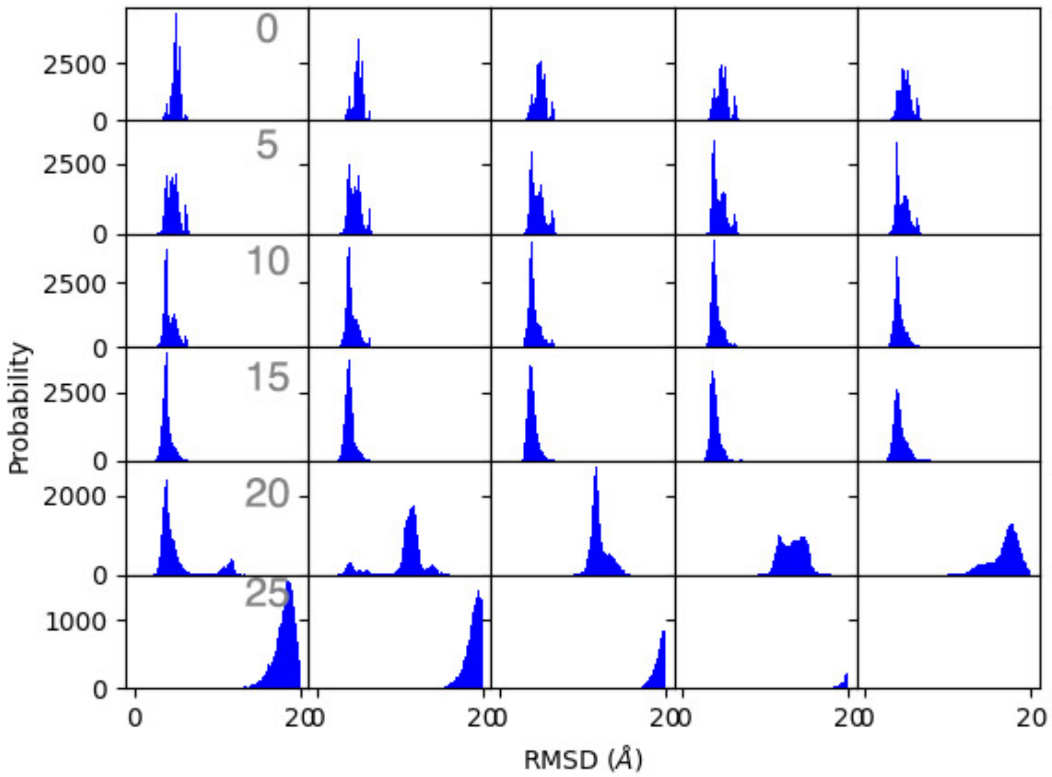

**Figure S13.** Protein-DNA RMSD distribution for each replica in the Replica Exchange ladder for the TATA system. Replica number increases to the right. Low replica index maps to low temperature and strong enforcement of the data (bound/misbound states). High replica indexes map to high temperature and vanishing restraints, enabling sampling of unbound states. Surprisingly, the most native-like conformations are identified in replicas 11-20, which show the lowest RMSD to native – this state has low population at the lowest replicas.

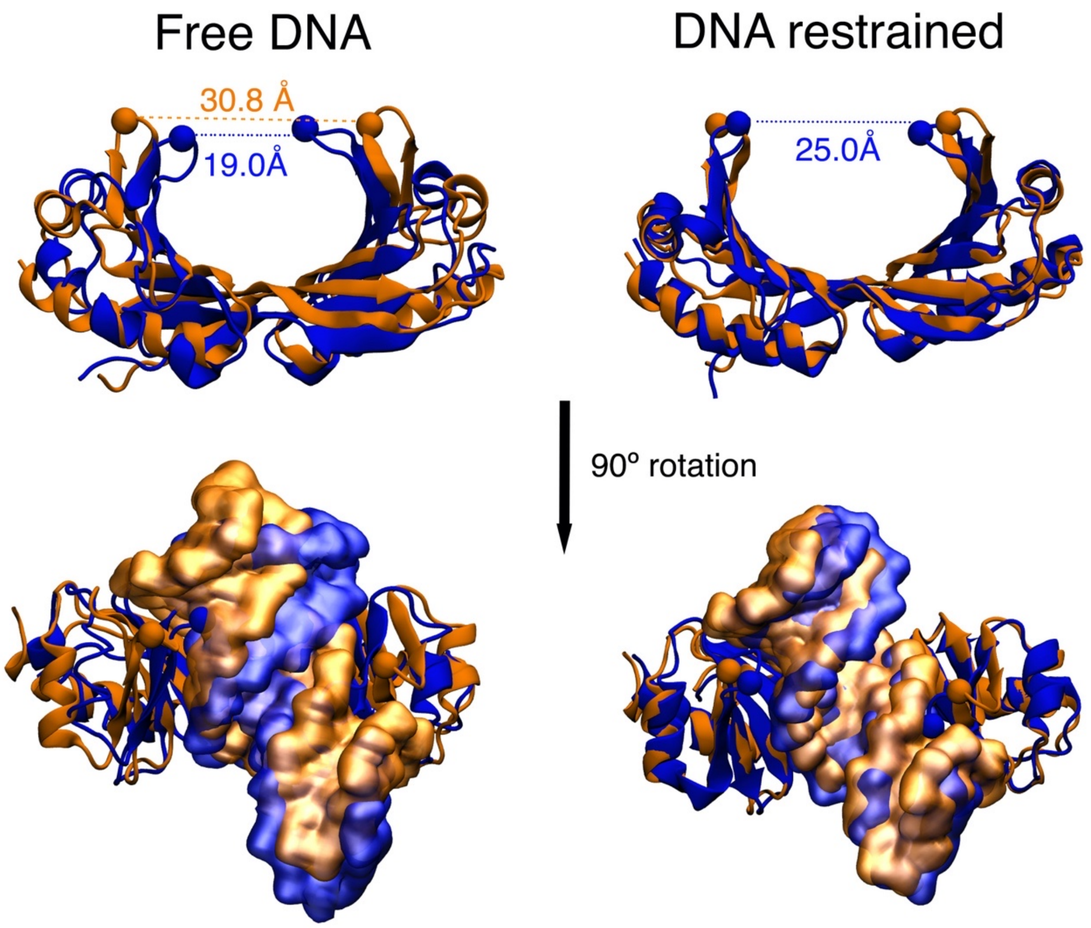

**Figure S14.** The TATA binding protein is highly flexible, modulating its ability to wrap around DNA's major groove. Native (orange) and simulated (blue) binding of the TATA box binding protein (cartoon representation) to DNA (space filling). In free simulations, the loop regions in the protein wrap more tightly around the DNA structure inducing a larger deformation (left). When the DNA is restrained with flat-bottom harmonic restraints, the protein still wraps around the major groove more tightly than in the experimental structure.

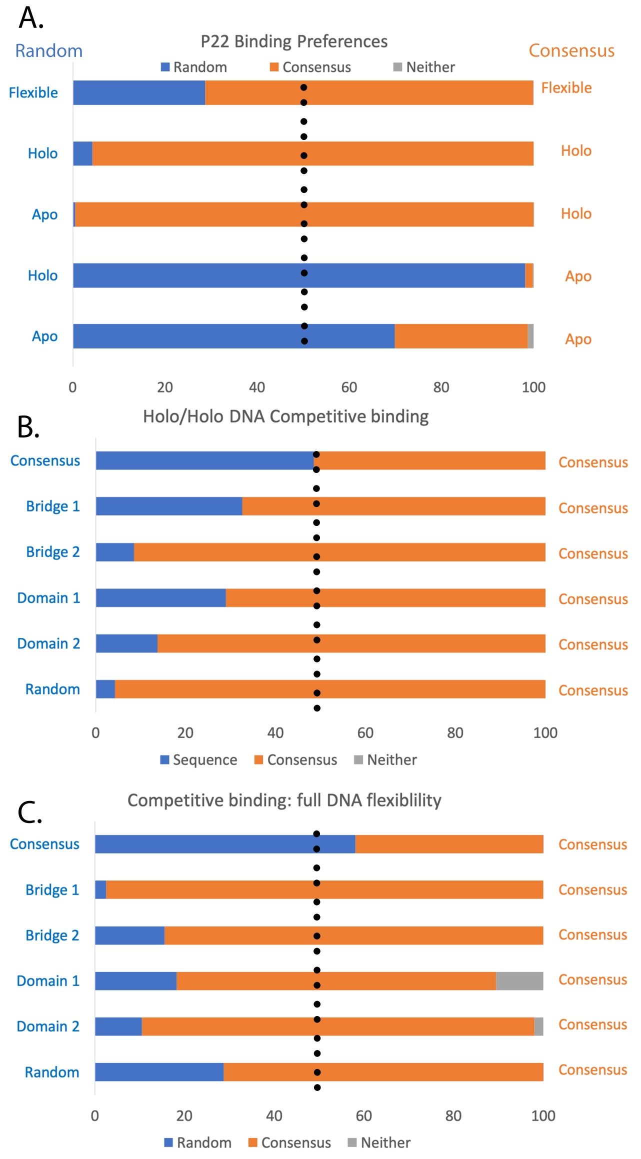

**Figure S15.** Competitive binding results for P22. Each bar represents a competitive binding simulation in which we compete a sequence/conformation (blue) against the consensus sequence in different conformations (orange). The bars represent a cumulative % one of the sequences is binding closer than 4Å from the correctly bound site. Conformations in which neither are found binding correctly are shown in grey. Panel A compares a random sequence with the consensus sequence when we place cartesian coordinate restraints to either the apo or holo conformation for each sequence. Results are also shown when both sequences are allowed flexibility to deform from apo to holo structures. Panel B compares 6 different sequences with consensus when all conformations are restrained to the holo conformation. Panel C shows all six sequences when full flexibility is allowed to each DNA sequence.
